# Brief Pulses of High-Level Fluid Shear Stress Enhance Metastatic Potential and Rapidly Alter the Metabolism of Cancer Cells

**DOI:** 10.1101/2025.04.09.647247

**Authors:** Amanda N. Pope, Devon L. Moose, Guy O. Hudson, Hank R. Weresh, Eric B. Taylor, Michael D. Henry

## Abstract

Circulating tumor cells (CTCs) face challenges to their survival including mechanical and oxidative stresses that are different from cancer cells in solid primary and metastatic tumors. The impact of adaptations to the fluid microenvironment of the circulation on the outcome of the metastatic cascade are not well understood. Here we find that cancer cells (PC-3, MDA-MB-231, Myc-CaP) exposed to brief pulses of high-level FSS exhibit enhanced invasiveness and anchorage-independent proliferation *in vitro* and enhanced metastatic colonization/tumor formation *in vivo*. Cancer cells exposed to FSS rapidly alter their metabolism in a manner that promotes survival by providing energy for cytoskeletal remodeling and contractility as well as reducing equivalents to counter oxidative stress associated with cell detachment. Thus, exposure to FSS may provide CTCs an unexpected survival benefit that promotes metastatic colonization.

## INTRODUCTION

Circulating tumor cells (CTCs) are intermediates in the formation of metastases at sites distant from primary tumors. They exist in the fluid microenvironment of the circulation, where they lack the supportive trophic factors of the primary tumor, adhesion to the extracellular matrix (ECM) and protection from the immune system [2]. They are also exposed to a range of hemodynamic forces [1], including fluid shear stress (FSS). Due to the dynamic nature of blood flow throughout the circulatory system, the levels of FSS a CTC may encounter vary greatly (from 1-6 dyn/cm^2^ in veins to >1000 dyn/cm^2^ in the heart; reviewed in [2]. Because most CTCs are larger than the average capillary diameter of ∼5 μm, they readily become entrapped after entering the microcirculation and may die, extravasate, or be dislodged to circulate freely again until they encounter the next capillary bed [3-5]. Considering blood flow velocity, the periods of free flow between longer periods of entrapment may be on the order of just seconds. Collectively, the death, destruction, and clearance of cells by extravasation or entrapment account for the relatively short (<2-hour) half-life of CTCs [6, 7]. Thus, although CTCs are exposed to a wide range of FSS, this exposure is brief and discontinuous.

How hemodynamic forces, including FSS, impact cancer cells is a growing area of interest [8]. It remains unclear if prior findings on adaptive responses in adherent cancer cells and non-transformed cells to exposure to FSS hold true for CTCs as these cancer cells exist in suspension. Compounding this uncertainty is that significant challenges remain in studying the effects of FSS on cancer cells, including improving the accuracy of *in vitro* modeling of the spatiotemporal dynamics of circulatory flow and *in vivo* tracking of the fates of individual CTCs [9]. Past studies of the effects of FSS on various cancer cell types used cancer cells in suspension to model CTCs and FSS of low levels (1-60 dynes/cm^2^) or long duration (15 minute-10 day). Thus, they did not account for either the upper end of the physiological range of FSS or the fact that CTCs are unlikely to flow continuously for such long periods. Nevertheless, they indicated that FSS can induce cellular phenotypes associated with metastasis, including increases in cell survival, adhesion, migration and invasion [10-13], expression of markers of EMT and stemness [14, 15], and tumorigenicity [15].

An additional barrier to survival encountered by CTCs is oxidative stress resulting from cell detachment. Oncogenic signaling can promote survival of detached epithelial cells by enhancing glucose uptake and flux through the pentose phosphate pathway to generate NADPH that combats detachment-induced oxidative stress [16]. Melanoma cells exhibit dependence on the folate pathway to generate NADPH and promote distant metastasis [17]. Indeed, recent work has revealed a variety of metabolic adaptations that CTCs may engage to survive reactive-oxygen species (ROS) toxicity [18]. The relationship between mechanical and oxidative stresses in CTCs is less clear. Exposure of MDA-MB-231 breast cancer cells to FSS (5-30 dynes/cm^2^) in a continuous flow closed loop model for up to 6 h increased ROS in a manner that was linked to enhanced cell migration [11]. Thus, ROS may either promote or prevent metastatic behaviors depending on the cellular and or microenvironmental/mechanical context.

In our previous studies, we found that cancer cells can resist mechanical destruction from brief, high-level exposure to FSS through engagement of a RhoA-dependent mechano-adaptive process resulting in cortical actin remodeling and actomyosin contractility that protects cells from mechanical damage [1]. Brief, high-level FSS exposure rapidly activates RhoA, consistent with its activation by a variety of mechanical stimuli. Because RhoA is involved in other cellular phenotypes that are associated with metastasis, such as invasion and survival in anchorage-independent conditions, we hypothesized that exposure to FSS in our model may not only protect cancer cells from destruction by FSS, but it may simultaneously promote metastatic behaviors in those surviving cells. Here we sought to determine the effects of brief exposure to higher levels of FSS that CTCs might encounter in circulation on metastatic behavior.

## RESULTS

### Exposure to FSS enhances metastatic phenotypes *in vitro* and *in vivo*

We exposed several cancer cell lines (PC-3, human prostate cancer, Myc-CaP, mouse prostate cancer, MDA-MB-231 human breast cancer) to millisecond pulses of high-level (937-1234 dyne/cm^2^) FSS in cell suspension over ∼10 minutes [19] and evaluated cellular phenotypes associated with metastasis as compared to cell suspensions held under static conditions for a similar duration. Exposure to FSS increased invasion in a Boyden chamber assay (**Figs. 1A-C**) and increased cell proliferation under anchorage-independent conditions (**Figs. 1D-F**) for all 3 cell lines. Since we previously demonstrated that in PC-3 cells exposure to FSS activates RhoA and RhoC, but not Rac1 [1], we determined if the increased invasive potential was dependent on RhoA or RhoC. We showed that in PC-3 cells the increased invasive potential conferred by FSS was dependent on RhoA, not RhoC (**Fig. S1A**).

**Figure 1.**
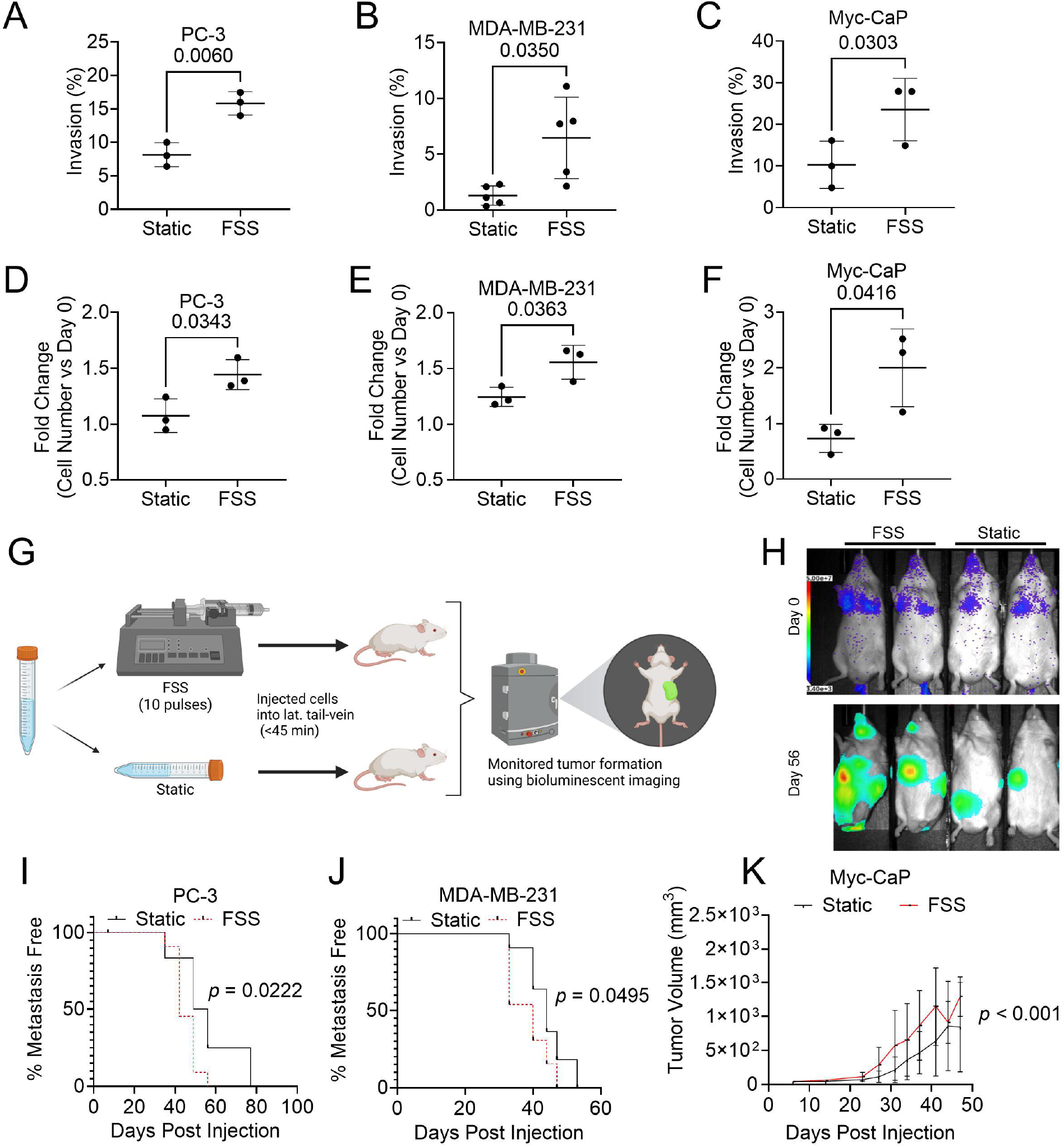
Exposure to FSS enhances metastatic potential *in vitro* and *in vivo*. (A-C) Effects of FSS exposure on invasion through a type I collagen matrix by PC-3 (A), MDA-MB-231 (B), and Myc-CaP (C) cells, 18 h post exposure (paired T-test; mean ± SEM). Effects of FSS exposure on number of PC-3 (D), MDA-MB-231 (E), Myc-CaP (F) cells at 24 h after seeding on poly-HEMA coated wells, expressed as fold change relative to initial cell number (paired T-test; mean ± SEM). (G) Schematic representation of *in vivo* experimental design made with *Biorender*. (H) Representative image of pairs of mice injected with either FSS exposed or static cells, on day of injection (Day 0) and 56 days after injection (Day 56). Color scale at Day 0 is the same for all images. Time to metastasis, with event time defined as BLI signal >10^7^ for (I) PC-3 and (J) MDA-MB-231(log-rank test, *p =* 0.0222, *p =* 0.0495, respectively). Mice inoculated with (K) Myc-CaP tumor volume measured by calipers (two-way ANOVA; *p =* 0.0069).

To test whether exposure to FSS of this same magnitude and duration can influence metastatic colonization or tumor growth, we exposed these cell lines to FSS and then injected equivalent numbers of viable, FSS- and static exposed cells into the appropriate animal hosts within 45min of FSS exposure and evaluated metastatic colonization using whole-body bioluminescence imaging (BLI) (**Fig. 1G,H**). In both PC-3 and MDA-MB-231 cells injected into the lateral tail vein, exposure to FSS decreased the time necessary to detect metastasis using BLI (**Figs. 1I,J; S1C,D**). More rapid development of metastatic colonies following FSS exposure is consistent with more cells surviving their journey through the circulation, potentially because prior FSS exposure increased their resistance to destruction in the circulation (as we showed previously [20]) and/or enhanced their ability to generate metastatic colonies as suggested above by the effects of FSS on cellular phenotypes. To address the former possibility, for PC-3 cells we found that there was no significant difference in the fraction of cells destroyed immediately following tail vein injection as compared static controls (**Figs. S1F,G**). We then evaluated the effects of FSS on cancer cells in immunocompetent mice. However tail vein injection of luciferase-expressing Myc-CaP cells in syngeneic FVB/NJ hosts resulted in aggressive metastatic colonization, with colonies that eventually regressed, likely due to an immune response to luciferase [21]. Exposure to FSS did not affect these findings (data not shown). As an alternative approach, we assessed subcutaneous tumor growth following exposure to FSS in parental Myc-CaP not expressing luciferase. Myc-CaP cells exposed to FSS exhibited faster tumor growth in syngeneic, immunocompetent hosts compared to static controls (**Fig. 1K, S1E**). Taken together, these findings indicate that exposure to brief pulses of high level FSS enhance metastatic phenotypes both *in vitro* and *in vivo*.

### Exposure to FSS alters the transcriptome and induces cytokine expression and oxidative metabolism

To investigate the mechanisms underlying the effect of FSS exposure on enhanced metastatic behaviors in cancer cells we exposed PC-3 cancer cells to the same protocol of FSS in the preceding studies and conducted RNA-Seq on FSS exposed and static samples 3, 12, and 24 h post FSS exposure. During this time course, cells were held in polyHEMA-coated dishes to simulate CTCs which are not attached to extracellular matrix (**Fig. 2A**). Principal component analysis revealed that the largest determinant of variation in gene expression overall is the time cells are held in suspension, but FSS exposure contributes to variation at any timepoint (**Fig S2A**). Over this time course, progressively more genes become significantly regulated (**Fig. 2B-D**). Gene Set Enrichment Analysis (GSEA) of the top regulated pathways revealed a significant increase in NF-κB-driven gene expression at 3 h which was not evident at later timepoints. At 12 and 24 h a gene expression pattern consistent with a proliferative response, including upregulation of myc, E2F targets, and other cell cycle-related genes emerged (**Fig 2F-G**), consistent with the FSS-induced anchorage-independent cell proliferation shown above (**Fig. 1D-F**). Additionally, at 12 h and 24 h post FSS exposure, it was notable that there was an increase in gene expression associated with oxidative metabolism including (*AOX1, COX6B2*, and *CTH*) (**Fig. S3A**).

**Figure 2.**
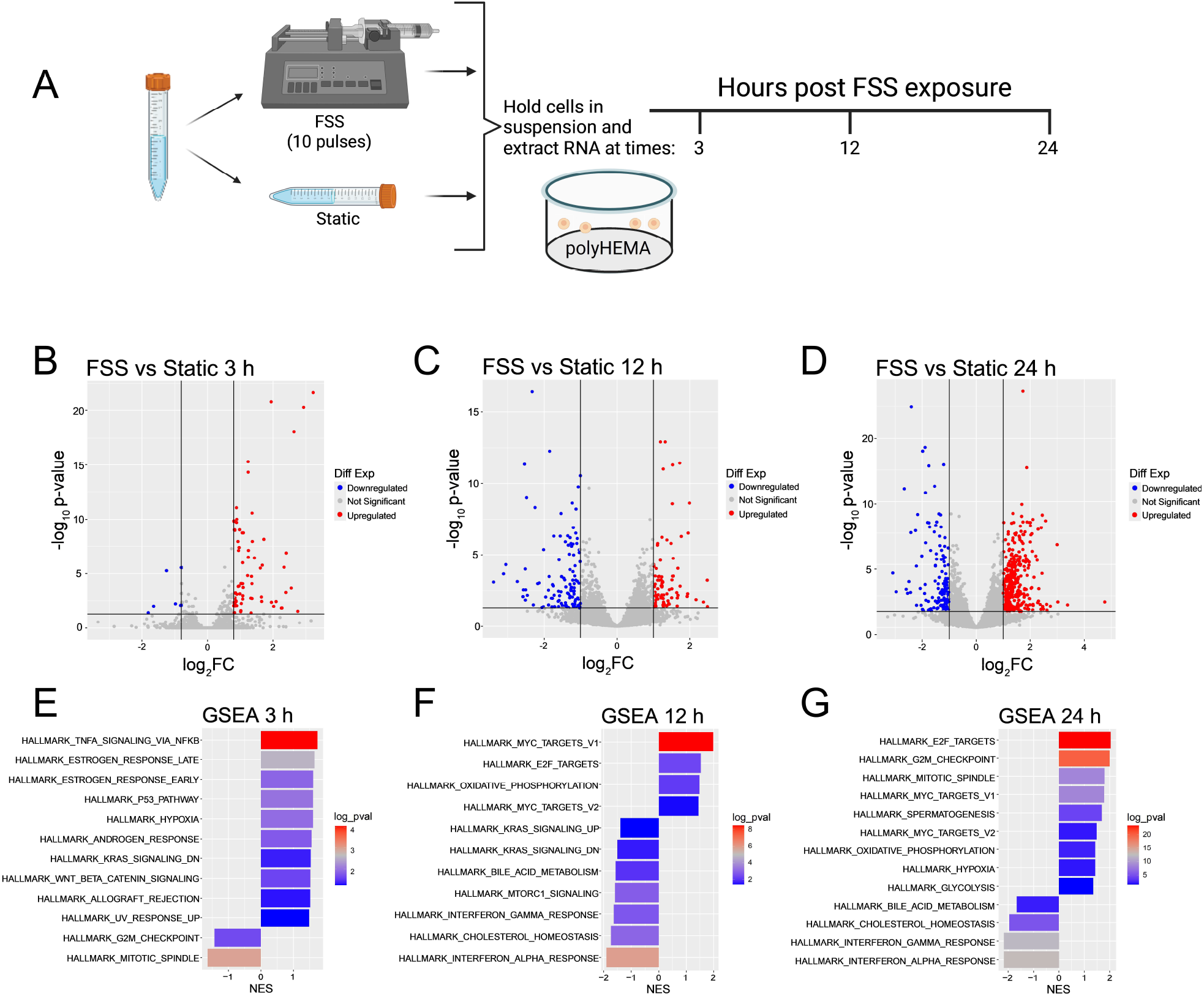
Exposure to FSS alters the transcriptome (A) Schematic representation of experiment using *Biorender*. (B-D) Volcano plots demonstrating up- or downregulated gene transcripts across three timepoints post FSS exposure: 3, 12, or 24 h, compared to static controls. (E-F) Gene set enrichment analysis (GSEA) across all three timepoints of significantly regulated pathways post FSS exposure compared to static controls.

To validate these findings, we conducted an NF-κB reporter assay in PC-3 and MDA-MB-231 cells. This showed that while NF-κB activity increases under both static and FSS exposed conditions from 0-6 h, consistent with a stress response in an attachment-deprived state [22], FSS exposure results in significantly higher NF-κB activity in both cell lines (**Fig. 3A,B**). We next evaluated the effect of FSS exposure on secretion of a subset of cytokines controlled by NF-κB. In PC-3 cells we found that FSS exposure significantly increased IL-6 and IL-11, and in MDA-MB-231 cells we found increased IL-6 and TGF-α within 24 h post exposure to FSS (**Fig. 3C,D**). Again, these cells were held in polyHEMA-coated dishes to deprive them of attachment.

**Figure 3.**
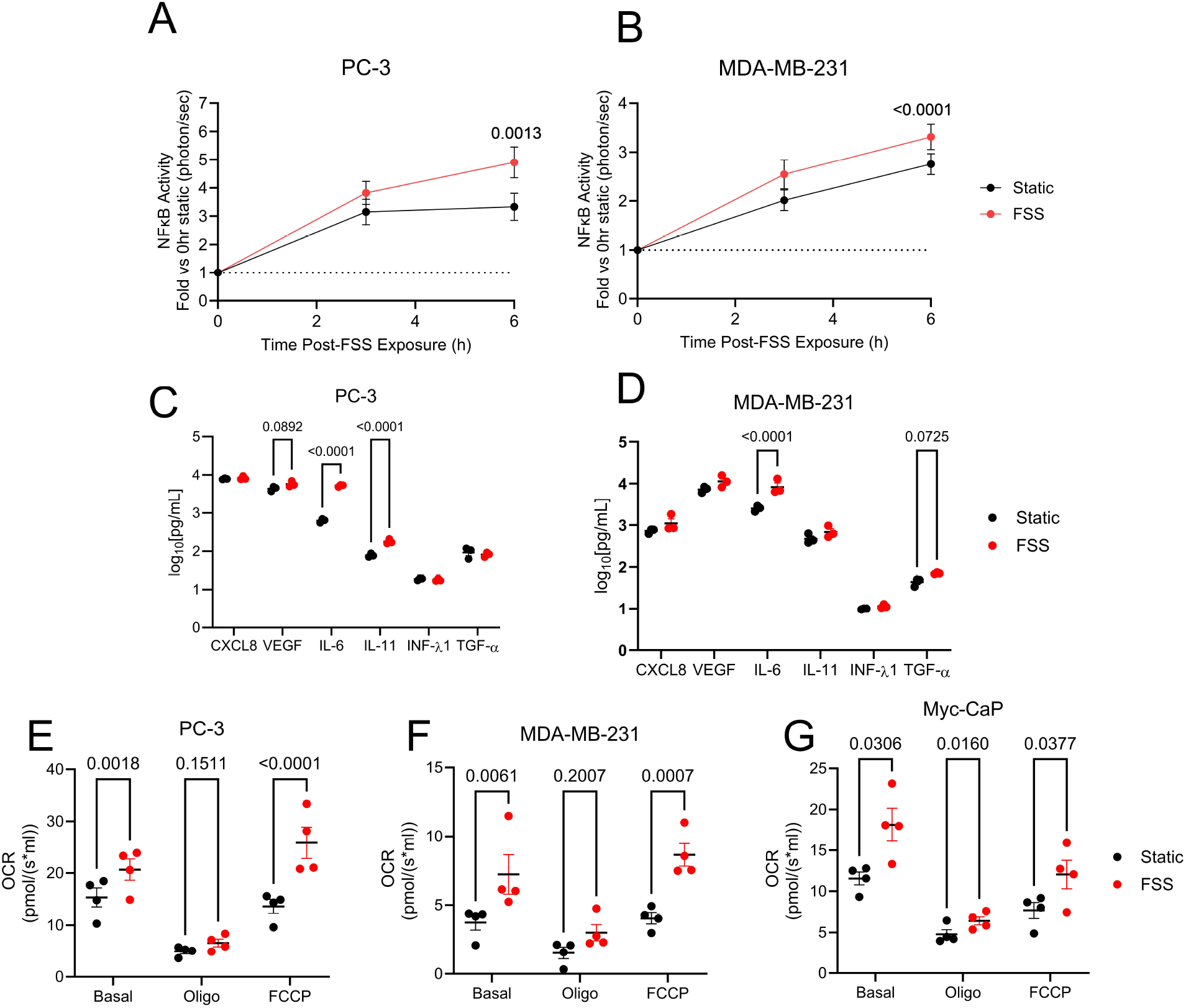
FSS exposure induces cytokine expression and oxidative metabolism. (A, B) Effects of FSS exposure activation of NFκB in PC-3 (A) and MDA-MB-231 (B). (C, D) Abundance of various cytokines in conditioned media from static and FSS exposed PC-3 (C) and MDA-MB-231 (D) (paired 2-way ANVOA, n=3, mean ± SEM). (E-F) Oxygen consumption rates (OCR) at basal, ATP-linked (0.0025 μM oligomycin), and maximum respiration (0.0025 μM FCCP) for PC-3 (E), MDA-MB-231 (F), and Myc-CaP (G) (2-way ANVOA; *n* = 4, mean ± SEM).

To assess the effects of FSS on oxidative metabolism we measured oxygen consumption in suspended cells 24 h post FSS (**Fig. 3E-G**). We observed a significant increase in basal respiration for all three cell lines. Treatment of cells with oligomycin revealed that there was no difference in ATP-linked respiration or proton leak while treatment with FCCP showed that FSS exposure increased maximal respiration in all 3 cell lines. Thus, within 24 h of exposure to FSS, cancer cells respond by increasing oxidative metabolism, consistent with upregulation of genes involved in oxidative metabolism evident in the transcriptomic analysis.

### Exposure to FSS rapidly alters the metabolome and engages folate metabolism to reduce ROS

Since numerous studies indicate that alterations in cellular metabolism underlie productive metastasis [17, 23-25] and our analysis of FSS induced changes in the transcriptome revealed that genes involved in oxidative metabolism were highly regulated, we performed metabolomic profiling of PC-3 and MDA-MB-231 cells immediately after exposure to FSS. Following the 10^th^ pulse of FSS, or under static conditions held for the same duration, cells were immediately plunged into liquid N_2_ to preserve the FFS-responsive and control metabolomes. Numerous metabolites were altered in both cell lines (**Fig. 4A,B**). We noted that the energy charge as evidenced by AMP/ATP and GMP/GTP ratios trended downward in PC-3 and was significantly lower in MDA-MB-231 cells (**Fig. 4C,D**). This finding is consistent with the fact that FSS exposure rapidly induces mechano-adaptation involving actin remodeling and actomyosin contractility-both energetically-costly functions [26].

**Figure 4:**
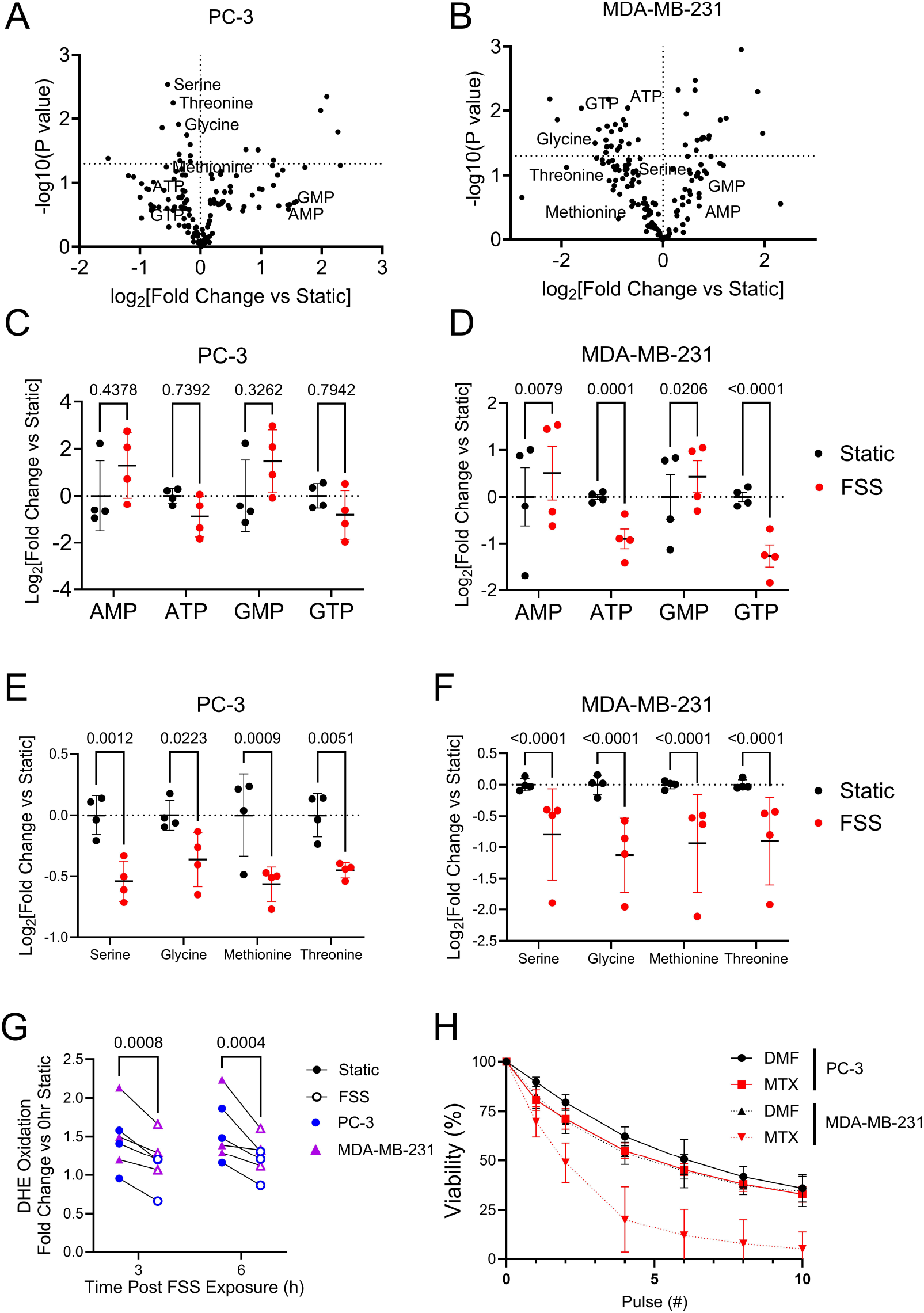
Exposure to FSS rapidly alters the metabolome and engages folate metabolism to reduce ROS. (A, B) Volcano plot from metabolomic profiling in (A) PC-3 and (B) MDA-MB-231 cells exposed to FSS (*n* = 4; paired t-test without correction). (C, D) Relative abundance of energy related metabolites in FSS-exposed and static PC-3 (C) and MDA-MB-231 (D) cells (*n* = 4; paired 2-way ANVOA with Bonferroni correction; *n* = 4, mean ± SD). (E-F) Fold change in relative abundance in metabolites that can feed the folate and 1-carbon cycle in PC-3 (E) and MDA-MB-231 (F) cell lines (*n* = 4; paired 2-way ANVOA with Bonferroni correction; *n* = 4 mean ± SD). (G) The effect of FSS exposure (open) relative to static controls (filled) on dihydroethidium (DHE) oxidation in MDA-MB-231 (magenta) and PC-3 (blue) (paired 2-way ANOVA, Bonferroni correction; *n* = 3, mean ± SD). (H) Effect of MTX (1µM, 4hr) on FSS resistance for PC-3 (solid line) and MDA-MB-231 (dashed line) cells (*n* = 3 for all conditions; mean ± SD).

To determine how cell metabolism might adapt to increased energy expenditure following FSS, we focused on the relative abundance of metabolites of glycolysis and the TCA cycle. There was not a consistent profile of changes in glycolytic intermediates across both PC-3 and MDA-MB-231 cell lines, although in PC-3 cells there were elevations in DHAP, 3-phosphoglycerate and lactate that are suggestive of increased glycolysis (**Fig. S4A,B**). We independently measured glucose uptake and lactate production immediately following exposure to FSS in PC-3, MDA-MB-231 and MyC-CAP cells. There were no significant differences in glucose uptake or lactate production between FSS exposed and static cells in any of these cell lines (**Fig. S4C-F**). Analysis of TCA cycle intermediates immediately following exposure to FSS revealed that there were no consistent significant similarities between PC-3 and MDA-MB-231 cells (**Fig. S5A,B**). While we did not find consistent evidence for immediate changes in glycolysis or TCA intermediates in response to FSS, we did find that treatment of PC-3 and MDA-MB-231 cells with 2-deozyglucose (2-DG) for 2 h sensitized both cell lines to destruction by FSS while it was not cytotoxic at this exposure (**Fig. S4G,H**). Taken together, these data indicate that while FSS exposure does not acutely alter glycolytic or TCA intermediates, glycolysis may be necessary to prepare cancer cells for exposure to FSS.

We noted that a subset of amino acids, serine, glycine, methionine and threonine, were consistently reduced in both cell lines immediately after exposure to FSS as compared to static conditions (**Fig. 4E,F**). These amino acids are associated with folate metabolism [27]. The folate pathway generates the reductant NADPH which acts via glutathione to combat ROS. We did not observe any significant differences in redox couples (NAD+/NADH; NADP+/NADPH; GSH/GSSG) in either PC-3 or MDA-MB-231 cells immediately, 3 h, or 6 h post FSS exposure (**Fig. S6A-H**). However, we did find that exposure to FSS significantly reduced ROS levels in both PC-3 and MDA-MB-231 cells 3 h and 6 h post exposure to FSS when held in suspension, indicating that FSS exposure leads to decreased levels of ROS in response to cell detachment (**Fig. 4G**). Moreover, lipid peroxides were also significantly reduced in response to FSS in a RhoA dependent manner (**Fig. S7A**). Treatment of PC-3 and MDA-MB-231 cells with a non-toxic dose of the folate inhibitor methotrexate (MTX) (**Fig.S8A**) resulted in dramatic sensitization of MDA-MB-231 cells to FSS but had little effect on PC-3 cells (**Fig 4H**). We note that MDA-MB-231 cells express higher levels of dihydrofolate reductase (DHFR), the target of methotrexate (**Fig. S8B**). Collectively, these data indicate that exposure to FSS triggers an increase in folate pathway flux, and that in MDA-MB-231 cells this protects cells from rapid destruction by FSS in the short-term. Thus, FSS exposure may trigger changes that protect CTCs from oxidative insults during metastasis.

## DISCUSSION

The data presented here indicate that cancer cells exposed to brief pulses of high level FSS rapidly alter their metabolism and gene expression profiles in a manner that promotes their fitness for metastatic colonization. Thus, not only do cancer cells have mechanisms that act to protect them from damage by FSS [20], FSS can be a stimulus for metastatic behavior. Indeed, FSS has potent biological effects on many cell types, most notably endothelial cells, which are constantly exposed to FSS [28]. Laminar flow models have been employed to determine the effects of FSS on adherent cancer cells in 2D culture [29-33], generally at low-level FSS (0.05-5 dynes/cm^2^). However, the relevance of these findings to CTCs, which are detached, are not clear. Although prior studies indicated that prolonged (15 min-10 d) exposure to lower-level (1-60 dynes/cm^2^) FSS induced cellular phenotypes *in vitro* and suggested that this potentially promotes metastatic behavior [15, 34], only one previous study examined the effects of FSS exposure *in vivo* [35]. It tested the metastatic colonization potential of lung cancer cells that had been exposed to FSS in in suspension (continuous flow loop with maximum wall shear stress of 35-918 dyn/cm^2^) for 72 hours before intracardial injection. Control cells were cultured under standard two-dimensional culture conditions (i.e., attached) and metastatic colonization by the FSS exposed cells was higher. Limitations of that study include that CTCs are unlikely to circulate continuously over such a long period of time, and it did not account for the effects of depriving cells of attachment (which our gene expression data indicate are significant; **Fig. S2A**)). Comparison of cells exposed briefly to FSS to counterparts held in suspension shows that FSS enhances metastatic potential. Our *in vitro* model of FSS exposure also has limitations. One is that the brief FSS exposure is at a high intensity that CTCs probably encounter only under certain circumstances, such as in turbulent flows around heart valves[36, 37]. Thus, *in vivo* only a small fraction of CTCs would be expected to be exposed to FSS at the same level as in our model. However, such unusual FSS exposure could have important effects on their metastatic potential. Nonetheless, taken together these data indicate that a broad range of FSS exposures to cancer cells in suspension have important biological effects on those cells which may model the effects of hemodynamic forces on CTCs in the circulation.

The results from our multi-omic study demonstrate that exposure to brief pulses of high-level FSS can enhance metastatic potential by inducing dynamic changes, in both gene expression and metabolism, that protect cancer cells from immediate destruction by FSS while promoting their survival and proliferation well after FSS exposure. An early change noted in the transcriptomic profile is transient expression of an NF-kB gene signature at 3 h. This is accompanied by enhanced secretion of certain cytokines including IL-6. Interestingly, Szczerba and colleagues showed that in heterotypic clusters of CTCs and neutrophils, the latter provided inflammatory cytokines, including IL-6, and acted to maintain proliferation of the CTCs [38]. Our data suggest that single CTCs exposed to brief pulses of high level FSS may achieve similar stimulation in a cell autonomous fashion that can promote metastatic colonization. Interestingly, our RNA sequencing data also suggested 12 and 24 h after FSS exposure that there is a reduction in response to type-I and type-II interferons (**Fig 2 F,G**). This may result in reduced immunogenicity of CTCs and promote metastasis [39]). Moreover, the Massagué lab demonstrated in lung cancer cells that inhibition of STING, a known promoter of the production of type-I interferons and an interferon-regulated gene [40], in cancer cells lead to a reduction in cancer cell dormancy [41]. Collectively, our data suggests that FSS exposure results in the multifaceted alterations in cytokine production/response that may act through varying mechanisms to promote proliferation of cancer cells and metastatic progression.

We do not yet know whether all the transcriptomic and metabolic alterations described here depend solely on mechano-adaptation driven by the RhoA-actomyosin axis. RhoA is known to activate YAP/TAZ in response to mechanical stimuli [33]. We also find evidence for activation of a YAP/TAZ gene signature in cancer cells after exposure to brief pulses of high-level FSS (data not shown). Thus, a RhoA>YAP/TAZ axis may contribute to anchorage independent survival and proliferation. We found here that RhoA was required for FSS-induced invasion in PC-3 cells consistent with its role in cancer cell invasion (reviewed in [42]). We have not yet established a role for RhoA in FSS induced cytokine secretion or metabolic alterations. However, RhoA and other RhoGTPases are known to activate NF-κB [43, 44], and RhoA and NF-κB collaborate to regulate mitochondrial glutaminolysis in breast cancer cells [45].

Although the role of RhoA is not yet clear, we noted a consistent depletion of amino acids associated with the folate pathway in both PC-3 and MDA-MB-231 cells immediately following exposure to FSS. Cell detachment provokes oxidative stress, and in various cell types prolonged exposure to FSS while suspended leads to increases in levels of ROS [11, 12, 16]. During metastasis, the folate pathway protects cells from oxidative stress, and folate inhibitors can reduce both CTC numbers and metastatic burden [17]. Our data demonstrate that exposure to FSS reduces ROS of PC-3 and MDA-MB-231 cells 3 and 6 hours after exposure, and rapidly reduces the abundance of amino acids that are methyl donors to the folate pathway. These data are in contrast with a previous study showing that continuous, long-duration (6 h) exposure to low levels of FSS results in elevated ROS, however, CTCs are unlikely to be continuously circulating for this period of time [11, 46]. Surprisingly, we did not see changes in redox couples 3 and 6 h after FSS exposure, suggesting that production and utilization of reducing capacity is in balance.

At longer timepoints following exposure to FSS, we find that cells increase oxidative metabolism. Prior evidence supports a shift to oxidative metabolism, and reduced glycolysis in melanoma cells under the conditions of detachment-induced stress [47]. Additionally, emerging data suggest that metastatic cancer cells upregulate oxidative phosphorylation and depend upon it in establishing secondary lesions or as CTCs [48-50]. Our data demonstrates that exposure to hemodynamic FSS promote oxidative phosphorylation, which might be a factor in the upregulation of oxidative phosphorylation in metastatic lesions that others have observed. Herein, our results highlight potential routes of reprogramming the metabolome immediately upon FSS exposure, as well as the lasting effects FSS has on regulating the transcriptome and metabolic pathways, which in turn enhance metastatic potential. In summary, this study reveals new mechanisms by which cancer cells adapt to FSS exposure, and these might be targeted in therapies to reduce the productive metastatic colonization of CTCs.

## METHODS

### Cell Lines

PC-3, MDA-MB-231, and Myc-CaP cells were obtained from the American Type Culture Collection (ATCC). PC-3 cells were maintained in DMEM:F12 (11320-033; ThermoFisher Scientific) with 10% FBS (S11150; Atlanta Biologicals) and 1x NEAA (11140-050; ThermoFisher Scientific). Both MDA-MB-231 and Myc-CaP cells were maintained in DMEM (11965-092; ThermoFisher Scientific) with 10% FBS and 1x NEAA. All cells were modified to express firefly luciferase through retroviral transduction, as previously described [19]. PC-3 cells were transduced with a control (SHC016; SigmaAldrich), RhoA (TRCN0000047711, TRCN0000047712; SigmaAldrich), or RhoC (TRCN00000291516, TRCN000047864; SigmaAldrich) short hairpin RNA (shRNA) using lentiviral particles, as described previously [20, 51].

### Chemical Reagents

2-deoxyglucose (2-DG) (D8375, Sigma-Aldrich) was dissolved in serum-free DMEM and cells were treated with 25mM 2-DG DMEM or DMEM 2 h prior to FSS exposure. Methotrexate hydrate (MTX) (13960, Cayman Chemical) was dissolved in dimethylformamide (DMF), and cells were treated with 1µM MTX or 1:10,000 DMF for 4 h prior to FSS exposure.

### Exposure to Fluid Shear Stress

Cells suspended in DMEM were exposed to 10 pulses of FSS by passage through a 30 gauge ½ inch needle (305106, BD) attached to a 5 mL syringe (309646, BD) at 250 µL/second using a syringe pump (70-3005, Harvard Apparatus), as described previously [52, 53]. A modification from published protocols is that the concentration of cells used for the FSS exposure in these experiments was 2×10^6^. Additionally for RNA sequencing and cell proliferations experiments, cells were sheared and subsequently plated in DMEM supplemented with 1x insulin-transferrin-selenium (ITS) (41400-045; ThermoFisher Scientific) and 5 mg/mL of bovine serum albumin (BSA) (03117057001, Roche).

### Measurement of Cell Viability

PC-3 (5,000) or MDA-MB-231 (10,000) cells were seeded in 96-well plates, with duplicates for each drug concentration tested. The day after seeding, the medium was changed to growth medium containing the stated concentration of MTX or DMF. Cells were treated for either 4 or 24 h and the medium was changed to growth medium without MTX. After 24 h, cell viability was measured using the resazurin (R7017, Sigma-Aldrich) dye.

### Clonogenic Assay

Static control samples were collected from the same wells as the treated samples and held in suspension while the remainder of the sample was exposed to FSS. For each condition, cells were plated in triplicate in a 6-well dish, at 500 cells/well. Cells were plated in growth medium and allowed to grow for 10 (PC-3) or 14 (MDA-MB-231) days before plates were stained with crystal violet solution and colonies (defined as >50 cells) were counted. Data from triplicate cultures were averaged and then normalized to the average of the biological replicates for the untreated, static control.

### Model of Metastatic Colonization

All procedures were approved by the University of Iowa Animal Care and Use Committee (protocols 5121574 and 8111574). To determine whether FSS exposure affects metastatic colonization, we injected 1×10^6^ viable trypan-blue negative cells into the lateral tail veins of NOD-*Prkdc*^*em26Cd52*^*Il2rg*^*em26Cd22*^/NjuCrl mice (572, Charles River) within 45 minutes of FSS exposure. Because of the need to minimize the time between FSS exposure and injection into the tail vein, only 2-5 mice were injected in a round. Each was injected with either sheared or static control cells initially collected from 2 15-cm tissue culture dishes. The cycle of FSS exposure followed by injection was repeated until 12 mice representing each condition (sheared and static) had been injected. To adequately power the experiment (α=0.05 and β=0.20), 12 mice per condition were used, and these were from 3 independent rounds of FSS exposure. Power analysis was based on a pilot study performed in NOD-Cg-*Prkdc*^*scid*^ *Il2rg*^*tm1Wjl*^/Szj (NSG) mice, where the mean time to metastasis (whole-body bioluminescence imaging signal (see below) >10^7^ photons/second) was 71±20 and 94 days for the FSS-exposed and static groups, respectively (**Fig 1-SFig1**).

### Bioluminescence Imaging (BLI)

After mice were injected with cancer cells, tumor burden was monitored by weekly bioluminescence imaging (BLI). For BLI, we performed intraperitoneal injection of 150 mg/kg of D-luciferin (LUCK, GoldBio), followed 5 minutes later by the detection of bioluminescence using an AMI HTX (Spectral Instruments Imaging); exposure time was 5 minutes. The AMIView Imaging Software was used to select an ROI that included the whole body, for quantification of signal intensity. The threshold for classification as metastatic disease was a whole-body BLI signal of ≥10^7^ photons/second.

### Invasion Assay

Invasion assays were performed using 8-µm, 24-well transwell inserts (3428, Corning) coated with 40µL of 0.8mg/mL rat-tail collagen type I (354236, Corning). The collagen matrix was allowed to polymerize for 30 minutes at 37ºC prior to adding the cells. 100 µL of 1×10^6^ cells/mL in DMEM was plated onto the matrix and 600 µL DMEM with 20% FBS was used as the chemoattractant in the lower chamber. Invasion was measured 18 h after seeding by adding D-luciferin to the media in the bottom of the transwell and the transwell insert, for a final concentration of 3 mg/mL and measuring the BLI signal. The medium was then removed from the top of the transwell insert and the non-migratory cells were scraped off before the BLI signal was measured a second time. The ratio of the invaded to total BLI signal was then used to determine the percentage of invading cells. BLI signal was measured for 2 minutes starting 5 minutes after luciferin was added at which point photon emission is stabilized. Duplicate samples were evaluated, and the data are represented as the average of the duplicates.

### Anchorage-Independent Cell Proliferation

For FSS-exposed and static conditions, trypan-blue negative cells (5×10^5^ cells/mL) were plated on 24-well dishes coated with poly-2-hydroxyethyl methacrylate (polyHEMA; P3932, Sigma-Aldrich) to prevent cell attachment, in DMEM with 1x ITS and 5 mg/mL of BSA. The concentration of these cells was again measured 24 h after FSS exposure and normalized to the original to determine the fold change in growth.

### In Vivo Destruction Assay

Prior to injection the PC-3 cells were either held in suspension or exposed to 10 pulses of FSS. We measured the fraction of cells destroyed immediately after lateral tail-vein injection as previously described [20]. Briefly, we injected 5×10^5^ luciferase expressing PC-3 cells in 200µL of PBS from either condition into NCI BALB/cAnNcr (555, Charles River) mice and within 3 minutes collected 500µL of blood via cardiac draw. The blood was then centrifuged at 1,500g for 5 min to separate the plasma from the cellular components. From the blood plasma, 100µL aliquots were transferred into 96-well black bottom plate before adding 100µL of assay buffer (200mM Tris-HCl pH=7.8, 10mM MgCl_2_, 0.5mM CoA, 0.3mM ATP, and 0.3mg/mL luciferin). A standard curve was generated by lysing 5×10^5^ luciferase expressing PC-3 cells in 200µL of 1% Tween-20 in ddH_2_O and then adding 13.3, 6.7, 3.3, 1.3, or 0µL of lysed cells to 500µL of blood collected from mice that had not been injected. The spiked blood was then processed the same as the blood from injected mice. The number of cells destroyed was then determined by linear regression.

### Western Blotting

40 µg of protein was loaded onto 12% SDS-polyacrylamide gels and transferred to PVDF membranes. Membranes were blocked for 1 hour using blocking buffer (927-90001, LI-COR) before being probed with primary antibodies (RhoA, ARH04, Cytoskeleton, INC; RhoC, D40E4, Cell Signaling Technology; αTubulin, 12G10, Developmental Studies Hybridoma Bank) overnight. Membranes were washed 3x before being probed with secondary antibody (IRDye 800 Goat α rabbit, 926-32211, LI-COR; IRDye 680 Goat α mouse, 926-68070, LI-COR) and signal was assessed using Odyssey (LI-COR) system.

### Metabolomic Profiling

Metabolomic profiling was performed on cells that were processed and snap frozen (using liquid nitrogen) immediately after they were exposed to 10 pulses of FSS or held in suspension as described above. Specifically, on the final FSS pulse, the cells were transferred into 10 mL of ice-cold PBS, the number of viable cells was determined, and the cells were centrifuged at 500g for 3 minutes at 4°C. Cell debris was removed from pellets by resuspension in ice-cold PBS, 2×10^6^ cells were transferred into a new tube, the cells were pelleted again, and these washed cell pellets were snap frozen.

### RNA Isolation

RNA was isolated from PC-3 cells that had been exposed to 10 pulses of FSS or held in suspension and then plated at 1×10^6^ cells/mL in 6-well plates that had been coated with polyHEMA to prevent cell attachment. For RNA extraction, the cytosolic contents were isolated by lysis in RLN buffer (50mM Tris-HCl pH=8, 140m NaCl, 1.5mM MgCl_2_, 0.5% NP-40) for 5 minutes, follwed by centrifugation at 12,000g for 5 minutes. Trizol was then added to the supernatant and the Direct-zol RNA Purification Kit (R2050, ZYMO Research) was used according to the manufacturer’s guidelines.

### RNA Sequencing

Samples were collected at 3, 12, and 24 h after FSS exposure. RNA sequencing and library preparation were performed by Novogene Co. LTD. Paired-end 150-bp sequences were generated using the Illumina platform. Sequences were aligned, and fragments counted, using the STAR aligner [54]. Differential gene expression analysis of the dataset was performed using DEseq2 [55] in Rstudio. The differential gene expression output was used to make a pre-rank list (log2 fold change * -log10 pvalue) of genes. Gene set enrichment analysis was then done on that pre-ranked gene list using fast gene set enrichment analysis (ref: https://www.biorxiv.org/content/10.1101/060012v3).

### Glutathione Measurement

PC-3 and MDA-MB-231 cells were exposed to FSS or held in suspension as outline above and 400µL samples of static and sheared conditions were taken 3 and 6 h after FSS exposure. Cells were pelleted by centrifuging at 500g for 3 minutes and the supernatant was discarded before the cell pellets were flash frozen on liquid nitrogen. To measure glutathione levels, samples were first deproteinized by adding 3 volumes of 5% 5-Sulfosalicylic Acid Solution to cell pellets, frozen and thawed twice and incubated on ice for 5 minutes. Samples were then centrifuged for 10 minutes at 10,000g to remove the precipitated protein. Supernatant was collected in a new tube and used to determine total glutathione content as described previously [39, 40]. The measurement of GSH uses a kinetic assay in which catalytic amounts of GSH cause a continuous reduction of 5,5-dithiobis(2-nitrobenzoic acid) (DTNB) to TNB and the GSSG formed is recycled by glutathione reductase and NADPH so the GSSG present will also react to give a positive value in this reaction. The yellow color of 5-thio-2-nitrobenzoic acid (TNB) generated is measured spectrophotometrically at 412 nm. The rate at which color accumulates is proportional to the amount of total glutathione. Reduced and oxidized glutathione were distinguished by the addition of 10µl 2-VP mixed 1:1 (v:v) with ethanol to 50µl of sample. The assay uses a standard curve of reduced glutathione to determine the amount of glutathione in the biological sample. Glutathione levels were normalized to the protein content.

### NADP(H) Measurement

PC-3 and MDA-MB-231 cells were exposed to FSS or held in suspension as outline above and 400µL samples of static and sheared conditions were taken 3 and 6 h after FSS exposure. Cells were pelleted by centrifuging at 500g for 3 minutes and the supernatant was discarded before the cell pellets were flash frozen on liquid nitrogen. Cell pellets were extracted using 200µL of 1% (DTAB) (D8638, Sigma) in 0.2N NaOH before adding 200µL of PBS. The measurement from the extracted cells was then performed using the NADPH/NADP Glo-Assay (Promega) following manufactures instructions. Luminescence was measured using AMI X imager with Aura software (Spectral Instruments). The concentration of NADP and NADPH were then converted to the total amount and normalized to protein concentration.

### Reactive Oxygen Species (ROS) Measurement

For both PC-3 and MDA-MB-231 cell lines, cells were subjected to either 10 pulses of FSS at 2×10^6^ cells/mL or held in suspension in phenol-red free RPMI with 1x ITS and 5mg/mL of BSA media. Cells were then plated on polyHEMA coated 24-well dishes at 5×10^5^ cells/mL in the phenol-red free RPMI with ITS and BSA. ROS were measured by incubating 1mL of cells with 10µM of dihydroethidium (DHE) for 30 minutes at 37C. ROS levels were measured immediately after FSS exposure, as well as 3 and 6 h later by flow cytometry using the data from 488nm excitation laser and comparing the ratio of the geometric mean to the initial static control.

### Lactate production

The rate of lactate production was measured using Lactate-GloTM assay kit (Promega, J5021). Our cancer cell lines were exposed to FSS or held in static conditions, washed with PBS, then immediately seeded on a 96-well plate in DMEM media. Lactate production was stopped using 5 μL of inactivation solution, provided by the kit, and placed on a plate-shaker for 3 min, followed by 5 μL of neutralization solution then on the plate-shaker for 1 min. 50 μL of lactate detection reagent was added then incubated for an 1 h at room temperature then luminescence was recorded. Lactate standard was diluted from 10 mM stock with a range from 0.1 mM to 0.00625 mM. The amount of lactate in each well was calculated from the standard curve. The rate of lactate production was calculated as:

Rate of lactate production = (lactate in cell culture)/ ((cell number well^-1^) x 24 × 60 × 60))

### Glucose uptake

The rate of glucose uptake was measured using Glucose-Glo™ assay kit (Promega, J6021). Our cancer cell lines were exposed to FSS or held in static conditions, washed with PBS, then immediately seeded on a 96-well plate in PBS. 50 μL of 2DG at 1mM was added to the samples then placed on a plate-shaker for 10 min. 25 μL of stopping buffer, provided by the kit, was added, followed by a quick shake, followed by 25 μL neutralization buffer added. 100µl of 2DG6P Detection Reagent was added then incubated for 1 h at room temperature. Luminescence was recorded. Glucose standard was diluted from 10 mM stock with a range starting at 0.1 mM to 0.00625 mM. The rate of glucose uptake was calculated as:

Rate of glucose consumption = (glucose in medium control-glucose in cell culture)/ ((cell number well^-1^) x 24 × 60 × 60)).

### Oxygen consumption rate (OCR) analysis

We exposed cell lines to FSS or held them in suspension and seeded them on polyHEMA coated plates for 24 h before measuring OCR. We used Oroboros Instruments Oxygraph-2k to measure basal, ATP-linked/proton leak, and maximal respiration at 7.5×10^5^ cells per replicate. ATP-link respiration/proton leak was induced by 0.0025 μM oligomycin and maximal respiration using 0.0025 μM FFCP. All measurements were recorded for 10 min intervals then averaged and reported in units of pmol min^-1^ cell number ^-1^.

### NFκB Activation Assay

PC-3 and MDA-MB-231 cells were transduced with lentiviral particles containing pHAGE NFκB-TA-LUC-UBC-GFP-W vector [45] (49343, AddGene, a gift from Darrell Kotton [56]) and flow sorted for GFP expression. To evaluate NFκB activation, cells were suspended in DMEM with 1x ITS and 5mg/mL BSA at 1×10^6^ cells/mL after exposing cells to FSS or hold them in suspension and 100µL was seeded in polyHEMA coated 96-well black bottom plate. Luciferase activity was assessed immediately and 6 and 24 h after seeding. Luciferase activity was normalized to the static 0 hour sample.

### Cytokine Measurement

PC-3 and MDA-MB-231 were held in suspension or exposed to FSS as outlined above and plated at 1×10^6^ cells/mL on polyHEMA coated 24-well. 24 h after FSS exposure, 500µL static and sheared samples were collected and centrifuged at 1500g for 15 minutes. The supernatant was collected and stored at -80C until the assay was performed. Cytokines were measured using a custom human 8-plex LEGENDplex panel (900001023, BioLegend). The assay was performed per manufactures instructions and flow cytometry on the beads was performed on a Cytek Aurora. Data analysis was performed using the LEGENDplex software from BioLegend. Data is presented as an average of duplicate reads for each biological replicate.

### Statistics

Within a cell line, differences in invasion by, and the proliferation of FSS-exposed vs static cells were analyzed for significance using the Students T-test. For the invasion data from the RhoA and RhoC knockdown experiments, the metabolomics data, and real-time PCR, repeated measures 2-way ANOVA with Bonferroni’s correction for multiple comparisons was used. Data for invasion are plotted as mean ± standard error of the mean; those shown in other graphs are plotted as mean ± standard deviation. Time to metastasis is presented as Kaplan-Meier curves and was analyzed using the log-rank test. The statistics described above were analyzed using GraphPad Prism 9. Principal component analysis was performed on metabolomics data using R v4.0.2. Differential expression analysis for RNAseq data within timepoints, as well as the effect size estimate for PCA analysis, were done using DEseq2 in R.

## Acknowledgments

We thank Eric Weatherford for insight and guidance for OCR analysis. We thank the staff and facilities of the Radiation and Free Radical Research Core, the Metabolic Phenotyping Core, the Metabolomics Core, and the Flow Cytometry Facility for their assistance and training. This work was supported by NIH award R01CA263550 to MDH. ANP was supported by T32 CA078586, DLM was supported by T32-GM0677954. Core facilities at the University of Iowa were supported in part by grant P30 CA086862 to the Holden Comprehensive Cancer Center. This research was also supported by a kind gift from the Sato Metastasis Research Fund.

## Author contributions

ANP, DLM, GFH, HRW investigation and validation; EBT data curation and formal analysis; ANP, DLM, MDH writing original draft; ANP and DLM visualization; MDH conceptualization, data analysis and funding acquisition.

## Figure Legends

Supplemental Figure 1: (A) Effects of knockdown of RhoA and RhoC on FSS-induced invasion (circles: shRNA construct 1; triangles: shRNA construct 2). Comparison of FSS-exposed vs static samples from each experiment was performed using a 2-way ANOVA with repeated measures and Bonferroni’s multiple comparison correction (shRNA target p=0.0360; FSS exposure p=0.0087; interaction p=0.0518; mean ± SEM). (B) Western blot showing RHOA, RHOC, and αTubulin levels in PC-3 cells expressing shSCR, shRHOA1, shRHOA2, shRHOC1, and shRHOC2. (C-E) Individual growth curves for each animal represented in Fig 1 I-K. (F) Schematic representation of *in vivo* cell destruction experiment using *Biorender*. (G) Percent of cells destroyed after injection into the lateral tail vein of mice, as measured by luciferase activity in blood plasma [20].

Supplemental Figure 2: (A) Principal component analysis of overall RNA sequencing data from FSS-exposed and static PC-3 cells held in suspension for 3, 12, or 24 h after FSS exposure.

Supplemental Figure 3: (A) Gene set enrichment plot for oxidative phosphorylation 24 h post FSS exposure, compared to static controls.

Supplemental Figure 4: (A, B) Effect of FSS exposure on the abundance of glycolytic metabolites in PC-3 (A) and MDA-MB-231 (B) cells (*n* = 4; paired 2-way ANVOA with Bonferroni correction; *n* = 4, mean ± SD). (C) Comparisons of glucose uptake between FSS-exposed and static cell lines, using Promega Glucose Uptake-Glo kit. (D) Calculated rates of glucose uptake using measurements from Figure 4 S3.C. (E) Comparison of lactate production between FSS-exposed and static cell lines, using Promega Lactate-Glo kit. (F) Calculated rates of lactate production using measurements from Figure 4 S2.E. (G) Effects of 2-deoxyglucose (2-DG) (25 mM, 2 h) on viability FSS resistance of both PC-3 and MDA-MB-231 cells following exposure to FSS (3-way ANOVA; 2-DG *p* = 0.0152, interaction 2-DG*FSS, *p* = 0.0003). (H) Effects of 2-DG (25mM, 2 h) on clonogenic potential of PC-3 (circle) and MDA-MB-231 (triangle) cells (Student’s T-test). (C-D) Statistical analysis consisted of paired t-test between respective cell lines; *n* = 4, mean ± SEM.

Supplemental Figure 5: (A, B) Relative abundance of TCA cycle metabolites from metabolomic profiling screen in FSS-exposed and static PC-3 (A) and MDA-MB-231 (B) cells (*n* = 4; paired 2-way ANVOA with Bonferroni correction; *n* = 4, mean ± SD).

Supplemental Figure 6: (A, B) Relative abundance of redox couple metabolites from metabolomic profiling screen in FSS-exposed and static PC-3 (A) and MDA-MB-231 (B) cells (*n* = 4; paired 2-way ANVOA with Bonferroni correction; *n* = 4, mean ± SD). (C, D) Ratios of redox couples from A and B from above to demonstrate redox status post-FSS exposure in PC-3 (C) and MDA-MB-231 (D) cell lines. (E, F) Effect of FSS 3 and 6 h after FSS exposure on NADPH/NADP^+^ ratios in PC-3 (E) or MDA-MB-231 (F) cell lines. (G, H) Effect of FSS 3 and 6 h after FSS exposure on reduced glutathione (GSH) oxidized (GSSG) ratios in PC-3 (G) and MDA-MB-231 (H) cells. All graphs are presented as mean ± SEM with each biological replicate being done in duplicate.

Supplemental Figure 7: (A) Measurements of lipid peroxides using Biodipy-C11 with flowcytometric analysis in PC-3 RhoA knockdown cells (shRhoA) or with non-targeting vector (shNT) (2-way ANOVA; *n* = 3, mean ± SD).

Supplemental Figure 8: (A) Dose-response of methotrexate (MTX) for acute cytotoxity (24 h after drug addition) in PC-3 (black) and MDA-MB-231 (red) cells with 4 hour (dashed line) and 24 h (solid line) of drug treatment compared to DMF control (*n* = 3 per cell line; mean ± SD). (B) Heat map for genes associated with folate pathway from CCLE database gene expression analysis.

