## Supplementary figures and images for "Brief Pulses of High-Level Fluid Shear Stress Enhance Metastatic Potential and Rapidly Alter the Metabolism of Cancer Cells"

### Supplemental Figure 1

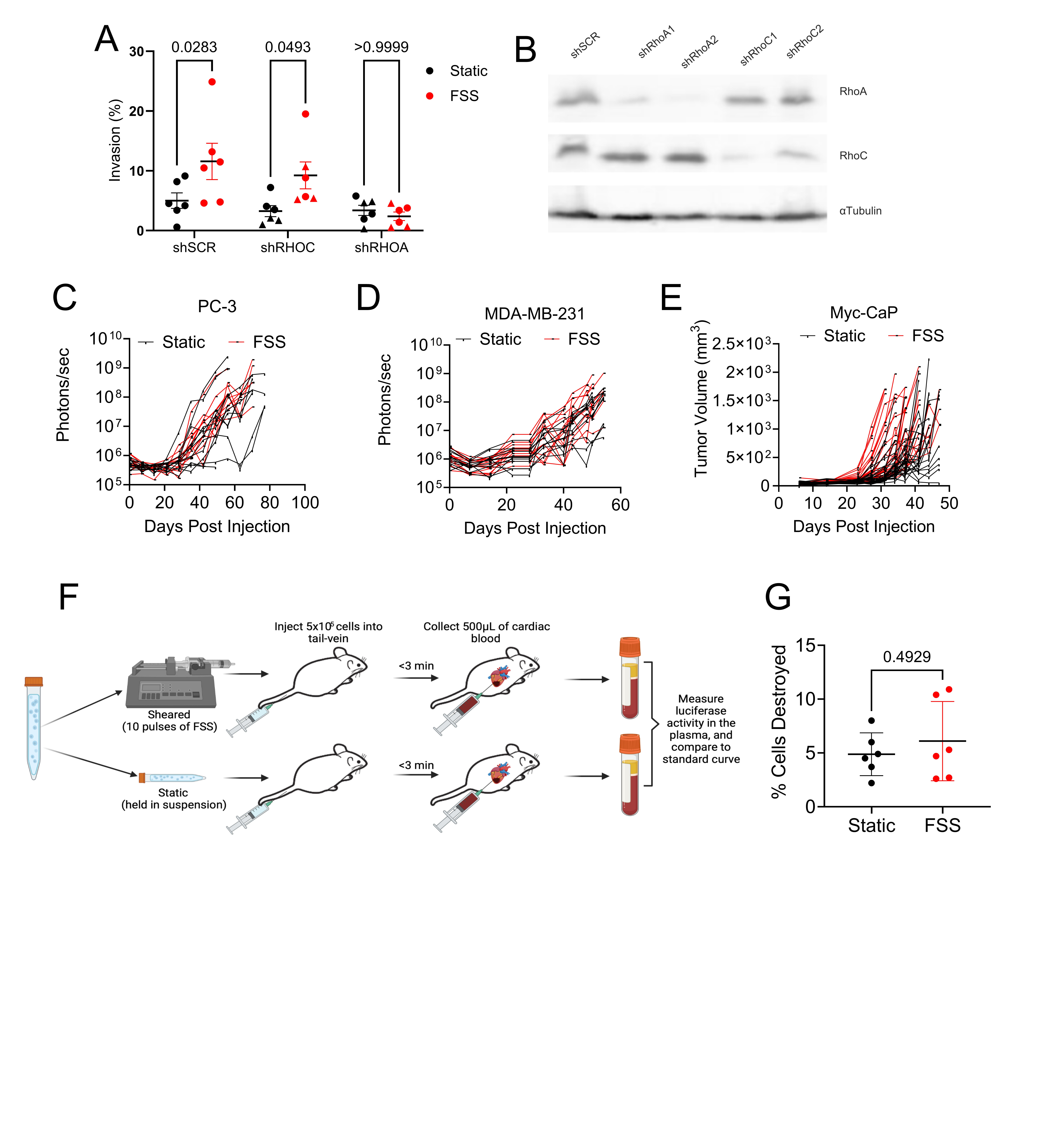

### Supplemental Figure 2

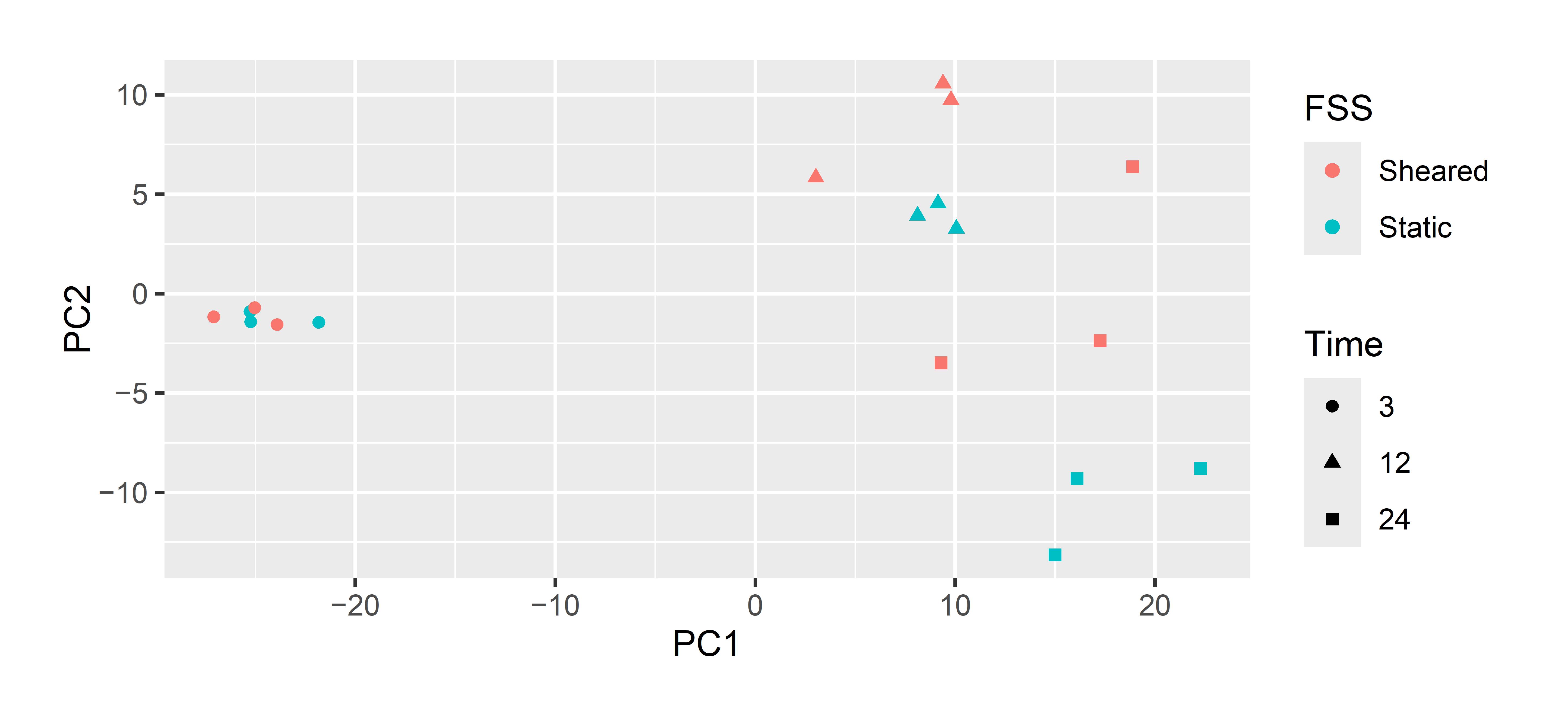

### Supplemental Figure 3

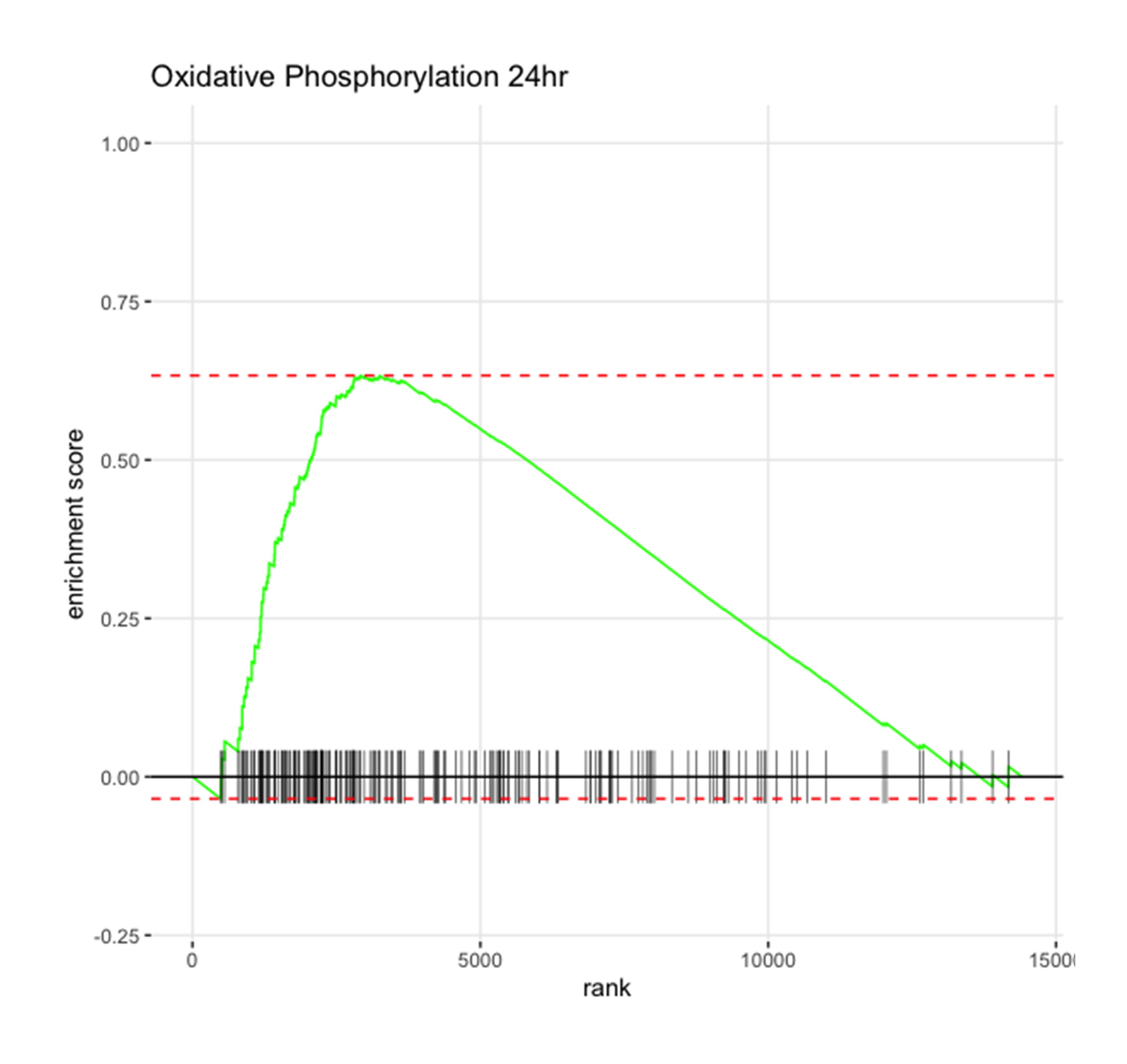

### Supplemental Figure 4

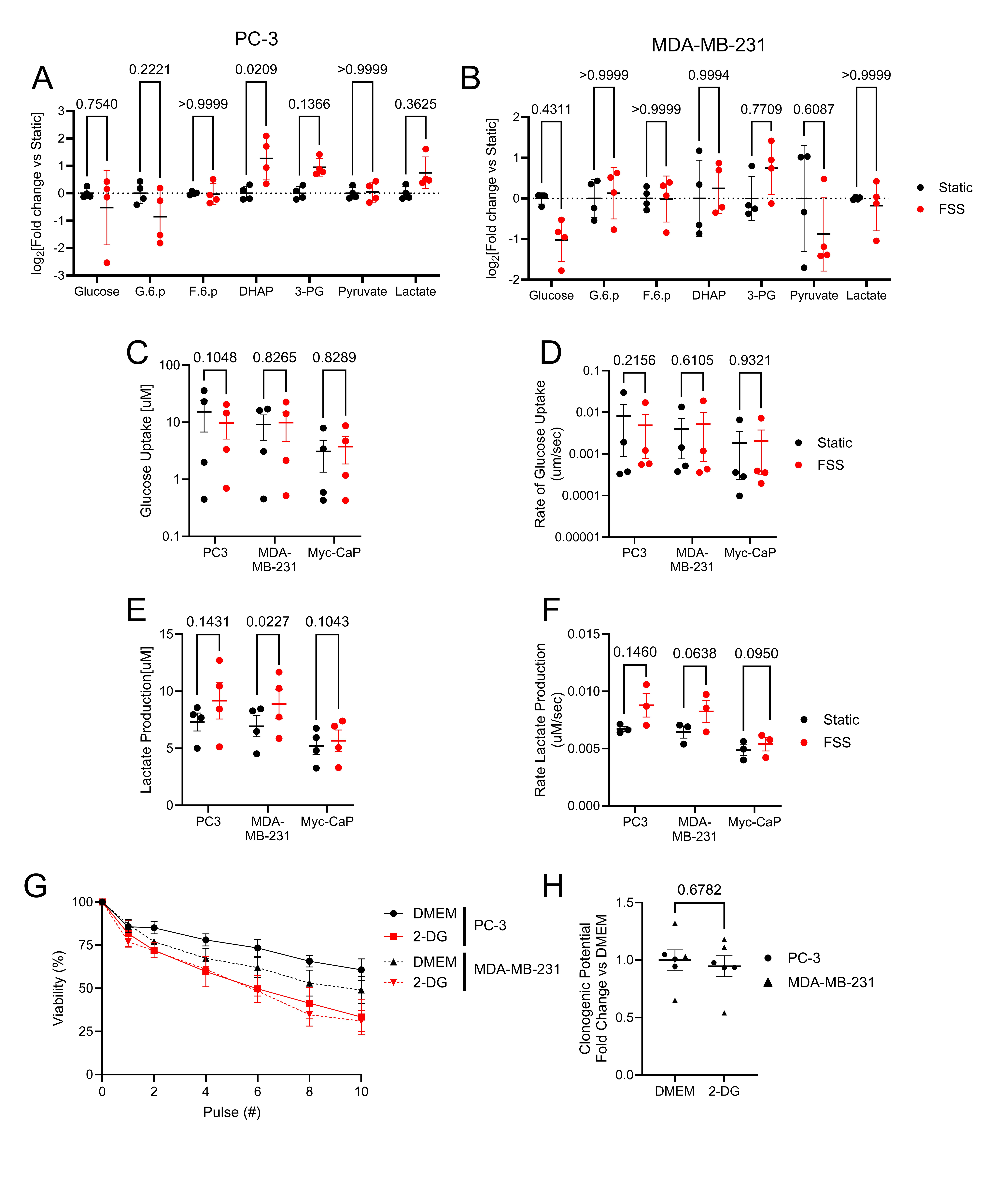

### Supplemental Figure 5

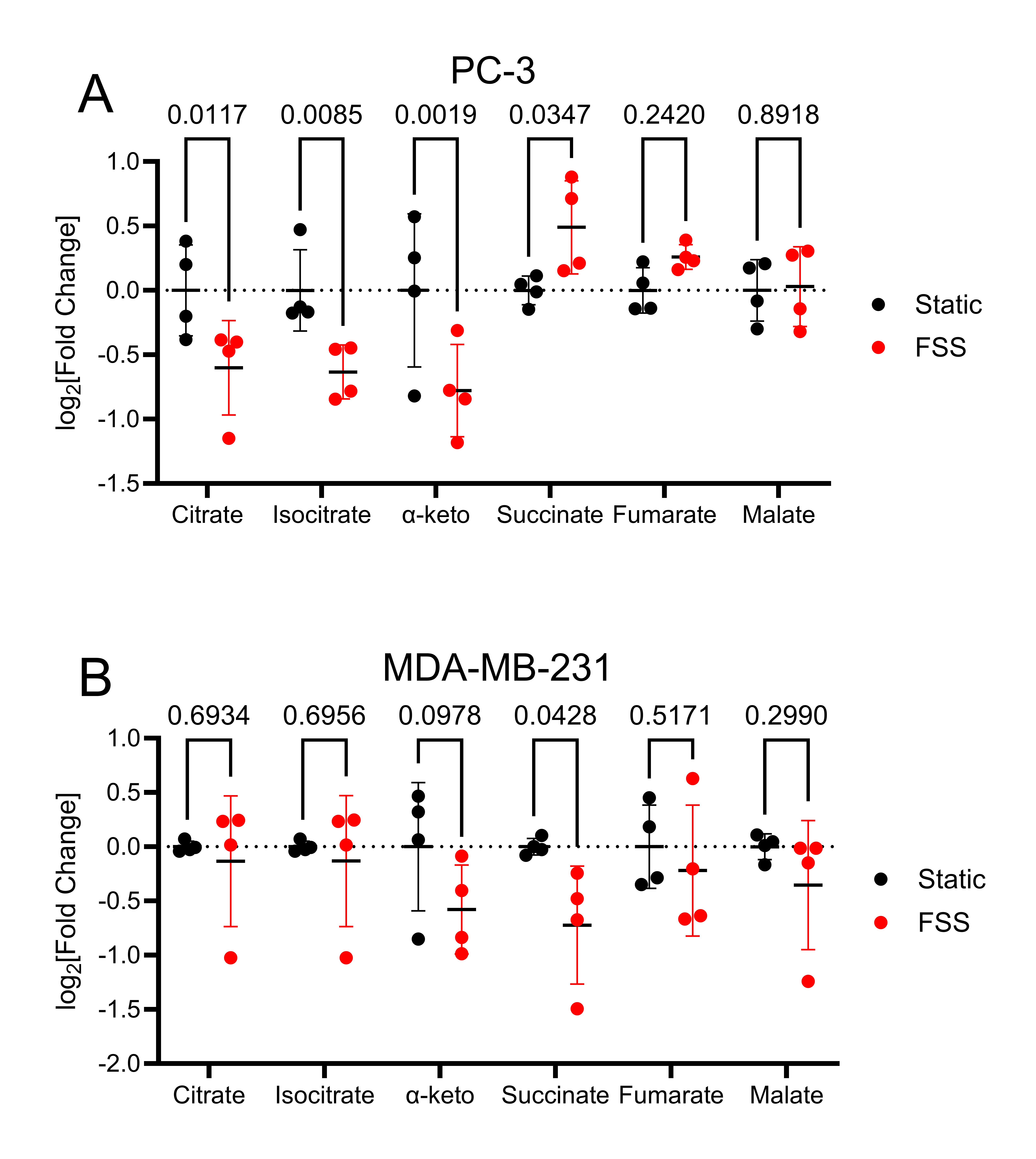

### Supplemental Figure 6

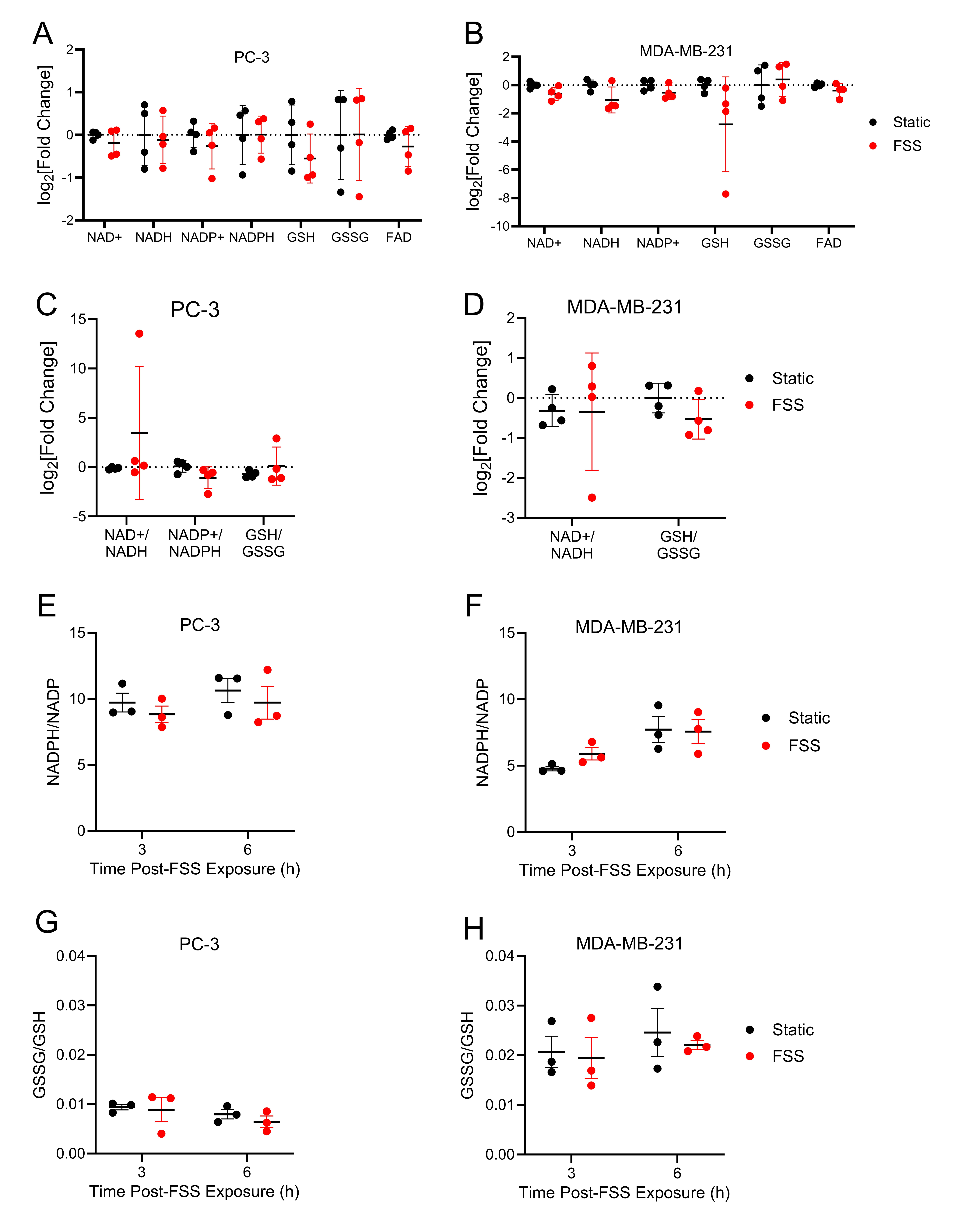

### Supplemental Figure 7

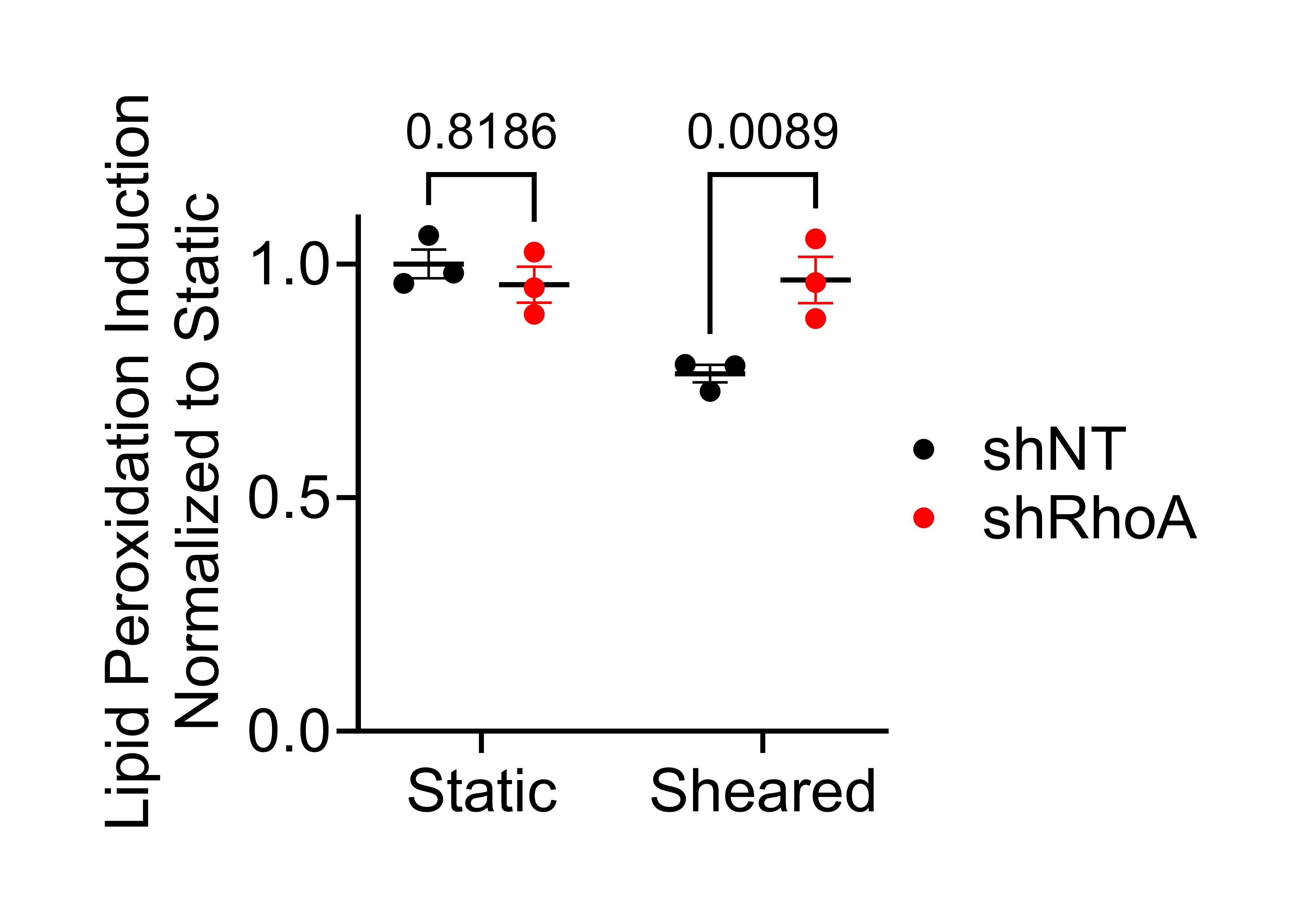

### Supplemental Figure 8

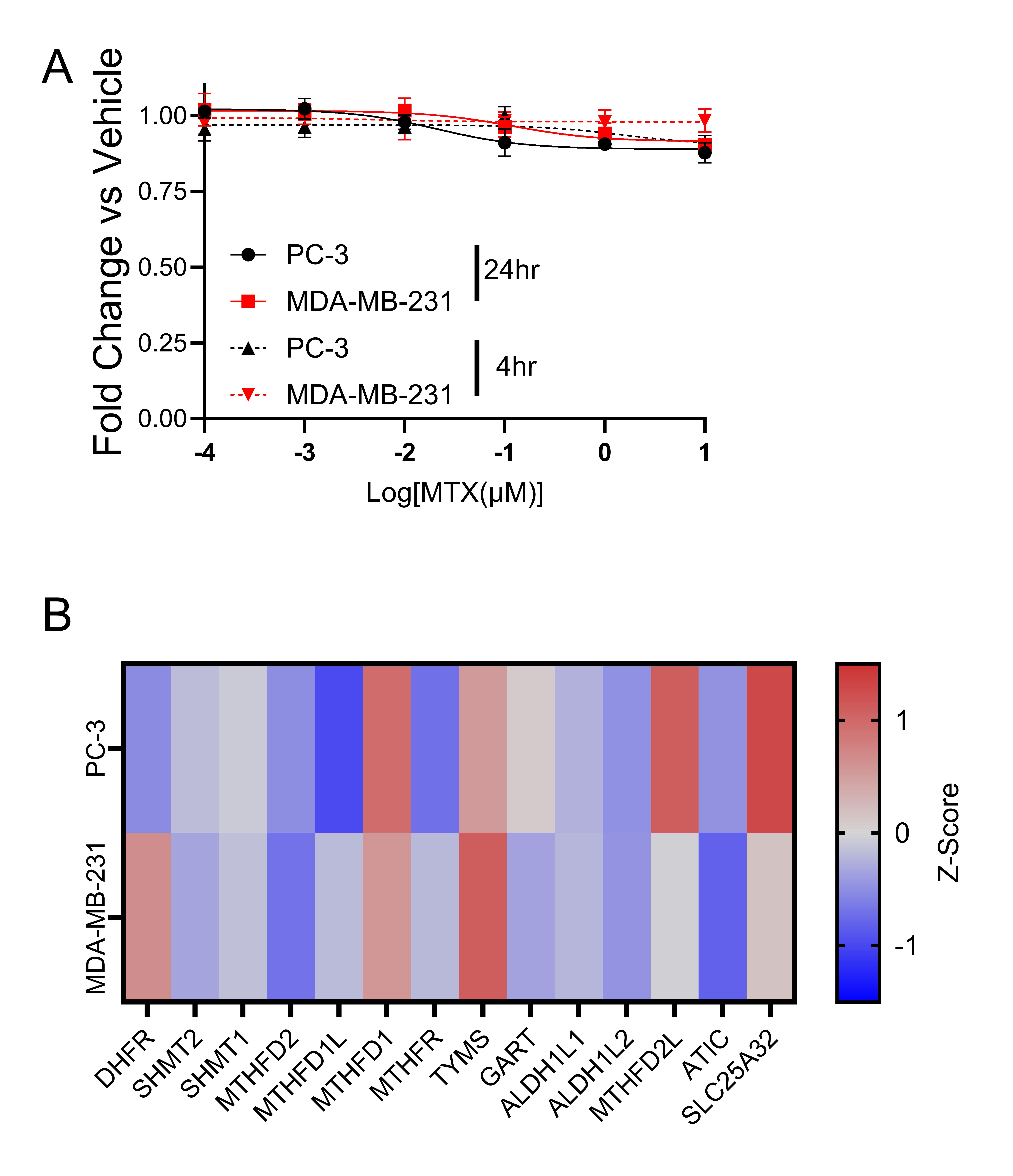
